# CHOPOFF: symbolic alignments enable fast and sensitive CRISPR off-target detection

**DOI:** 10.1101/2025.01.06.603201

**Authors:** Kornel Labun, Oline Rio, Håkon Tjeldnes, Michał Swirski, Anna Zofia Komisarczuk, Emma Haapaniemi, Eivind Valen

## Abstract

CRISPR/Cas systems offer powerful tools for genome editing, but their therapeutic application is hampered by the risk of unintended off-target mutations. Many molecular methods have been established to detect off-target editing, however, their sensitivity depends on first identifying potential sites using *in silico* methods. However, these *in silico* prediction methods are challenged by a trade-off between speed and sensitivity, and can fail to comprehensively detect all edited off-target sites. Here, we demonstrate that ignoring bulges has led to missing editing at off-target sites in previous studies and that continuing this practice can lead to inflated claims of fidelity. As a solution, we introduce the concept of symbolic alignments to efficiently identify off-targets without sacrificing sensitivity. We further present specialized data structures that enable rapid, alignment-free probabilistic ranking of guide RNAs based on their predicted off-target burden. Implemented in the tool CHOPOFF, these innovations accommodate mismatches, bulges (insertions/deletions), and incorporate genomic sequence variants for personalized off-target assessment. Benchmarking demonstrates that CHOPOFF significantly outperforms state-of-the-art tools in both prediction accuracy and computational speed.

**Availability:** CHOPOFF command line available at https://github.com/JokingHero/CHOPOFF.jl

CHOPOFF web server available at https://crisprtools.org/chopoff

## Main

CRISPR/Cas is a transformative genome editing technology that has dramatically improved our ability to introduce precise changes into genes and genomes. Due to its versatility and relative ease of use, CRISPR/Cas is widely employed in basic research, most notably functional genomics, disease modeling, synthetic biology, and many others. Additionally, given its capacity to edit genetic errors, CRISPR/Cas is also being evaluated for a broad range of medical applications, recently resulting in the regulatory approval of the first CRISPR-based therapeutics ^1^.

Despite its transformative potential, CRISPR/Cas technology has notable limitations and liabilities, the most important of which is the potential for off-target editing, whereby CRISPR/Cas cuts DNA at unintended locations. Such off-target (OT) edits can distort the interpretation of functional experiments and introduce noise and variability, thus decreasing the reliability of experimental results and functional conclusions. OT activity is especially dangerous in therapeutic applications of CRISPR, where even a very low-frequency OT edit may have disastrous outcomes ^2,3^. To address this challenge, many efforts in the field have focused on minimizing OTs. First, several methods to improve guideRNA (gRNA) design *in silico* have been developed to maximize target specificity ^4^. In parallel, molecular methods to measure OT effects after running an experiment have been developed to quantify and validate these predictions. These include GUIDE-seq ^5,6^, CIRCLE-seq ^7^, and SITE-seq ^8^, which have been instrumental in increasing our capacity to quantify real OT edits. A limitation of all these genome-wide molecular OT profiling assays, however, is that they also depend on *in silico* identification of potential OT sites before molecular quantification of their activity. Computational methods to predict potential OT sites are therefore vital to both pre-experiment prediction of potential OT targets and for validating the outcome of those experiments. Suboptimal methods, particularly at the validation phase, can therefore lead to severely underestimating the magnitude of OT editing. This can potentially give a false impression of ‘high-fidelity’, even when the approaches are ostensibly ‘experimentally verified’.

CRISPR/Cas OTs prediction algorithms work by conducting a genome-wide search to identify sites with high complementarity with the gRNA sequence and the presence of protospacer adjacent motif (PAM) sequence, which is essential for Cas cutting. Given that gRNA has been shown to edit sites with up to 6 mismatches and even with small bulges (insertions or deletions), predicting CRISPR OT effects represents a significant challenge ^5,9,10^. Identifying potential OT sites is therefore equivalent to finding all short sequences in a genome that are likely to engage with a given gRNA despite the presence of mismatches, insertions, and deletions. Currently, commonly used sgRNA design tools such as CHOPCHOP ^11^ and CRISPOR ^12^ restrict OT searches to 3-4 mismatches and do not support bulges. Importantly, many of the published papers when analyzing data from genome-wide OT profiling assays, ignore bulges due to the complexity they introduce^5,13–20^. Several tools that can handle mismatches and bulges, such as Cas-OFFinder ^21^, CRISPRitz ^22^, and SWOffinder ^23^, have been developed, but they are computationally intensive, and their running time makes these approaches impractical in larger experiments or as web tools. Additionally, support for natural genomic variation (SNPs, insertions, and deletions) in existing software is limited ^11,12,21,23^, which restricts the utility of CRISPR/Cas technology for precision medicine applications.

### Reanalysis of GUIDE-seq reveals off-targets with bulges

To demonstrate the consequences of ignoring bulges when searching for OTs, we re-analyzed the original GUIDE-seq data ^5^ with a method that allows for mismatches, insertions, and deletions (Levenshtein distance). The original analysis was restricted to only mismatches (Hamming distance) and a bulge only at the gRNA/PAM interface. Our re-analysis revealed several sites missed across the 6 guides in this data set. Examples of 4 sites across 2 guides (HEK site 1 and EMX1) are shown in **Figure 1A,D**, where deletions and insertions close to the PAM make the Hamming distance much larger than the Levenshtein distance, resulting in them being ignored in the original analysis. The patterns of editing, as represented by GUIDE-seq reads, at those sites (**Figure 1C,F**) are similar to editing at the on-target site (**Figure 1B,E**), albeit with an expected decrease in editing efficiency. As an independent validation, we performed a simple cell-free screening for 27 variable guides on the same on-target loci - where alignment between sgRNA and target will create variable sgRNA-OT conditions such as mismatches, bulges, and deletions (**Supplementary Figure 1A**). The outcome was measured by gel electrophoresis. This revealed that several deletions and insertions between sgRNA-OT are tolerated and create detectable cut rates (**Supplementary Figure 1B**).

**Figure 1.**
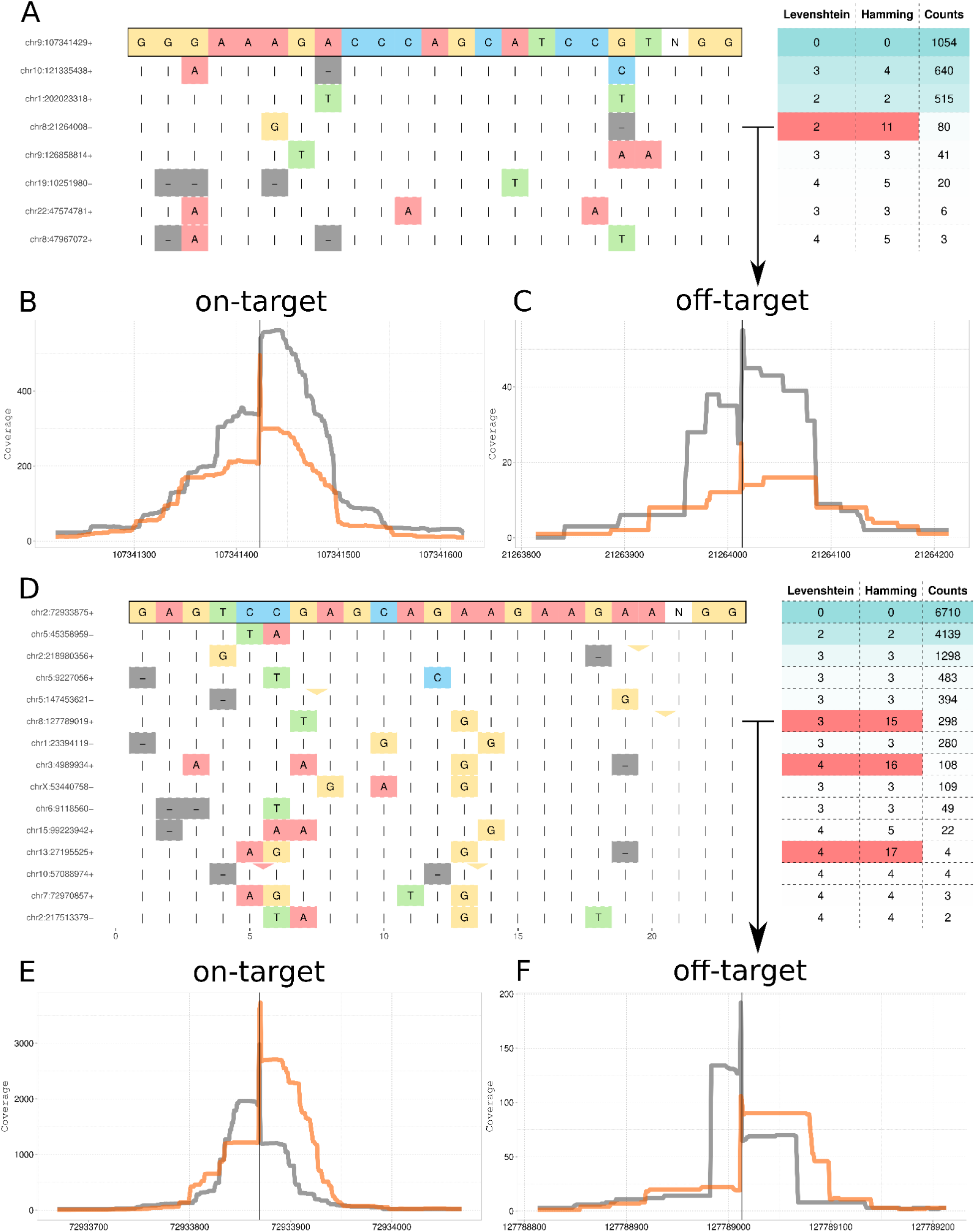
Off-target editing also occurs with insertions and deletions, and a bulge close to the PAM will render the site undetectable to mismatch-based tools. **A)** Re-analyzed data from original GUIDE-seq^5^ showing distance measured by Hamming (only mismatch) and Levenshtein (mismatch/insertions/deletions) for HEK site sgRNA 1. We marked with red the sites that will evade detection when searching only up to 6 mismatches. Deletions are displayed as grey dashes, while insertions are represented by triangles with color indicating the inserted base. **B)** Coverage view of the on-target site for HEK site sgRNA 1. Orange and black lines are for the sense and antisense GUIDE-seq read coverage. **C)** Previously undetected site for off-target site for HEK site sgRNA 1 due to deletion close to the PAM. **D)** Re-analyzed data for EMX1. **E)** Coverage view of the on-target site for EMX1. **F)** Previously undetected site for off-target site for EMX1 due to insertion of G close to the PAM.

### Levenshtein distance is a superior metric to Hamming distance for OT prediction

To better understand the relationship between editing efficiency and the two distance metrics, we re-analyzed off-target datasets from three different studies: GUIDE-seq ^5,6^, dual-target ^24^, SURRO-seq ^17^. Dual-target and SURRO-seq datasets consist of artificial synthetic oligo sets of sgRNA-OT pairs, while GUIDE-seq data is an analysis of actual genomic OTs. Current studies, including the data sources, typically simplify the relationship between sgRNA and OT sites by only considering Hamming distance. This, however, ignores sites that are highly similar but have misaligned regions due to an insertion or deletion. We therefore realigned the sgRNA and OT sites using the Levenshtein distance as a metric and contrasted this with the OT Hamming distances. The OT editing efficiency was calculated as relative to the on-target efficiency. For the majority of OT loci, the Hamming and Levenshtein metrics are identical (**Supplementary Figure 3**), and these loci (GUIDE-seq: 1033, dual-target: 3314, SURRO-seq: 2654) therefore represent the expected editing efficiencies at a given Hamming distance. We then asked for the OTs where Levenshtein is lower than the Hamming distance, which metric more accurately represented the OT site, i.e. are these OT sites edited more than their Hamming distance would predict? In total, 510 OT sites fulfilled this criterion (GUIDE-seq: 229, dual-target: 155, SURRO-seq: 126), and we found that regardless of experimental methodology, Levenshtein distance consistently gives a better estimate of the editing efficiency (**Supplementary Figure 3**).

### Computational predictions at greater distances produce many false positives

From the reanalysis of these three datasets, one can also estimate how the tradeoff between sensitivity and specificity for OT detection changes with increasing distance to the on-target. All datasets agree that there is a substantial loss in editing efficiency with increasing distance (**Supplementary Figure 3**). When averaged across all OT sites, both Hamming distance and Levenshtein distance of 3 are at the edge of detection for all 3 methods. There is, however, a large variance across guides and a few outliers. For instance, there exist outliers with a distance of 4 with relative editing rates above 0.5. However, these are almost no longer found for distances above 4. Larger and more comprehensive datasets can, in the future, give a more comprehensive overview and demonstrate whether these observations hold in general.

These estimates are based on the editing efficiency of OTs relative to the on-target efficiency (“relative efficiency”), so we wanted to investigate whether the conclusion holds when considering absolute editing efficiency. It has previously been argued that GUIDE-seq reads can not be used directly to estimate editing efficiency due to variable efficiency of dsODN tag integration, influence of cell-specific DNA repair pathways, and PCR amplification biases ^6,25,26^. Despite this, GUIDE-seq data is highly correlated with PCR-based methods of OT validation, as demonstrated in the original paper ^5^ and later confirmed ^6^. We replicated this relationship in our analysis (**Supplementary Figure 2A**, R = 0.67, p < 2.2e-16), but also observed that normalizing the GUIDE-seq data by the on-target editing efficiency (**Equation 1)** provides a good approximation for the true OT editing efficiencies as measured by independent methods (**Supplementary Figure 2B**, R = 0.87, p < 2.2e-16). This simple normalization allows repurposing GUIDE-seq reads from those studies where independent methods have evaluated the on-target efficiency and constructing a more comprehensive dataset of estimated OT editing efficiencies. We found that applying similar normalization for CHANGE-seq did not yield similarly predictive results.

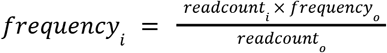

**Equation 1**. Inferring indel frequency on the off-target sites using GUIDE-seq reads. Where *frequency*_i_ is the editing efficiency at off-target i, *readcount*_i_ is the read count at off-target i, estimated by GUIDE-seq, *frequency*_0_ is the editing efficiency at on-target site, estimated by rhAmpSeq/PCR, and *readcount*_0_ is the read count at on-target site, as estimated by GUIDE-seq.

Based on the GUIDE-seq dataset from Lazzarotto et al. 2020, we used this relationship to summarize how many OT sites are edited and at what efficiency, in relation to all potential OT sites, for 51 sgRNAs **Table 1** ^6^. To combine results from many sgRNAs, we normalized GUIDE-seq counts to the on-target site as described above and calculated the mean editing efficiency. Multiplying the chance of a given OT being edited (**Table 1**, ‘mean fraction of edited sites’) by the relative editing efficiency for edited sites gives us the expected editing at any random OT site (**Table 1**).

**Table 1.** Summary of GUIDE-seq for 51 sgRNAs and their off-target. Mean editing efficiency is the editing efficiency of our re-scaled GUIDE-seq counts as in the equation above, averaged across all guides. After on-target normalization, this measure is averaged across many guides. Expected editing is the fraction of edited OT sites multiplied by the expected editing efficiency at a given edited OT site.

| Distance | Mean editing efficiency [%] | Number of edited OTs sites | Number of OTs sites within distance | Mean fraction of edited sites [%] | Expected editing [%] |
| --- | --- | --- | --- | --- | --- |
| 0 | 61.27 | 51 | 51 | 100.00 | 61.27 |
| 1 | 38.57 | 4 | 8 | 50.00 | 19.29 |
| 2 | 7.92 | 77 | 281 | 27.40 | 2.17 |
| 3 | 2.57 | 390 | 7117 | 5.48 | 0.14 |
| 4 | 0.78 | 468 | 129297 | 0.36 | 0.003 |

These estimates show that while the number of potential OT targets grows exponentially with Levenshtein distance, the probability of each site being cut decreases dramatically with increased distance (**Figure 2A, Table 1**). At a distance of 3, there are 7117 possible OTs of which approximately 5.48% will be edited at a relative efficiency of approximately 5.73%. This is equivalent to an expected editing of about 0.14% for any given OT site, and an expectation that about 94.52% of computationally predicted OT sites will be false positives. At a distance of 4, the expected editing is 0.0003% and out of 129297 possible OT sites, 99.64% will be a false positive. This leads to the situation where, at greater search distances, the vast majority of predictions based on the use of Levenshtein distance that exceed a certain threshold are false positives, i.e. sites that an algorithm would predict as “cuttable” but in practice are not going to be cut. Indeed, targets at an edit distance of 4 are extremely unlikely to be cut (0.36%), and if they are cut, it is typically at a negligible efficiency (0.78%, **Figure 2A, Table 1**).

**Figure 2.**
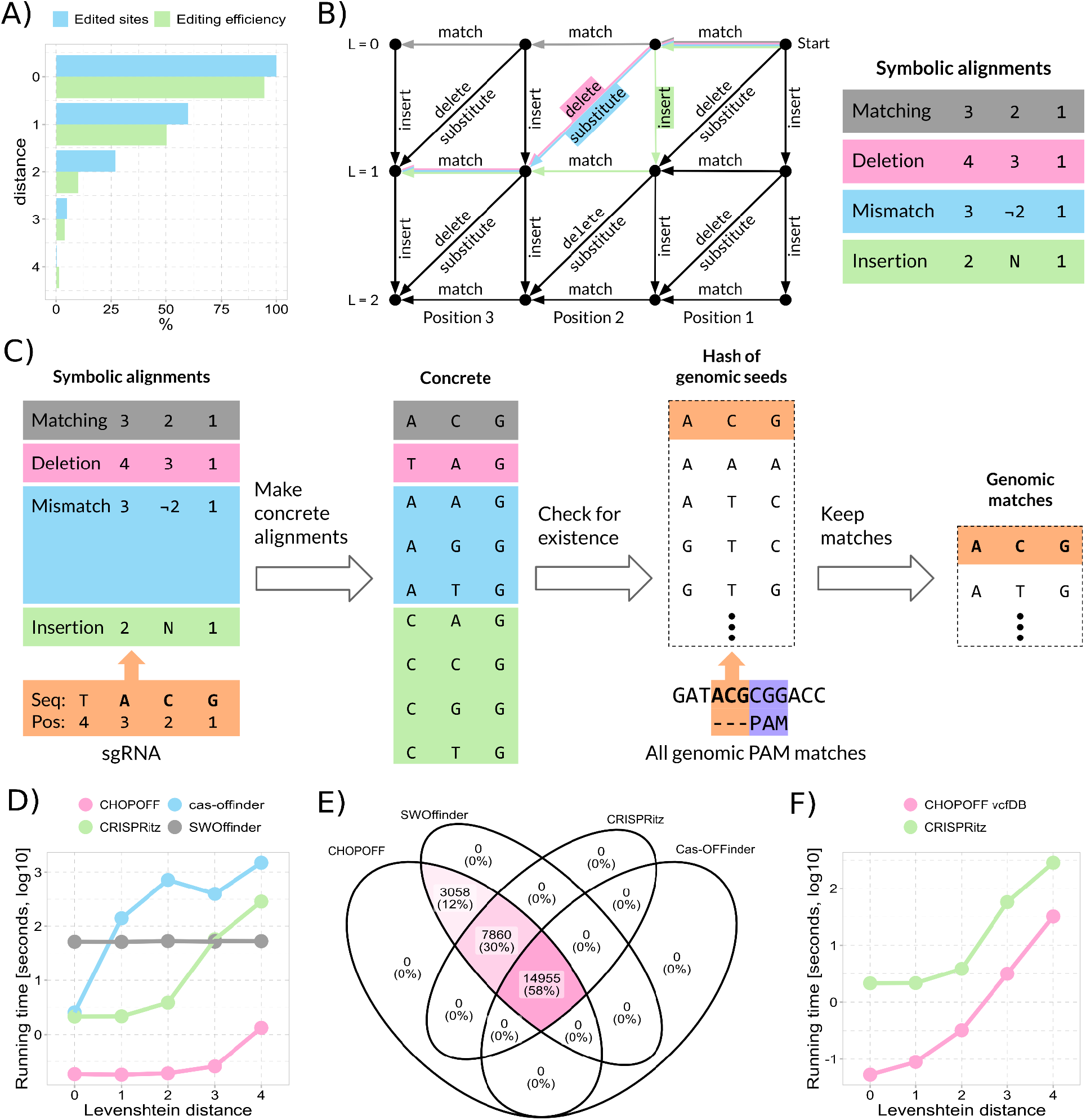
Precomputed symbolic alignments combined with hashing are fast and precise at finding all off-targets. **A)** GUIDE-seq data for 51 sgRNA shows the percentage of sites edited (blue) and the mean editing efficiency of those sites (green) with increasing Levenshtein distance (y-axis). **B)** All possible alignments for an sgRNA sequence can be pre-computed through the enumeration of all paths in a graph (left). The red path highlights the only zero distance (L=0) alignment, while orange, blue, and green paths show example alignments at distance one. The right panel shows examples of precomputed symbolic alignments corresponding to the paths highlighted in the graph. **C)** All symbolic alignment paths are pre-computed. When presented with a concrete sgRNA sequence, the symbolic alignments are substituted with the nucleotides of the sgRNA, giving a list of all possible off-target sequences for that sgRNA. These are checked against a genomic index, and if a match is found, a Needleman-Wunsch alignment is made between the sgRNA and the loci. **D)** Running time for methods supporting bulges with increasing Levenshtein distance. **E)** Venn diagram showcasing the sensitivity of OT search by each tool. **F)** Running time for methods supporting genomic variation.

### Symbolic alignments enable fast and sensitive CRISPR off-target detection

Taken together, this analysis suggests that an OT prediction algorithm should include the capacity to account for short bulges, in addition to mismatches, but also minimize false positives by not searching for sites that are too dissimilar. Aligning sgRNA sequences with a genome at an arbitrary Levenshtein distance is a time-consuming computational operation. However, since the length of the sgRNA and PAM positioning are known prior to the search, and given our conclusions regarding the optimal Levenshtein distance, we reasoned that precomputing all possible OT sequences in advance would allow us to identify OTs without the need to align all sgRNAs with the entire genome. We implemented this precomputing framework into CHOPOFF, a standalone software and framework for identifying potential OTs based on the idea of preprocessing both the sgRNA and the genome. CHOPOFF is an order of magnitude faster than comparable methods while supporting arbitrary Levenshtein distance, genomic variation and ambiguous bases through IUPAC codes.

The genome is preprocessed by identifying all potential target sites defined by being adjacent to a valid PAM sequence (∼230 * 10^6^ for SpCas9 in humans). These sites are stored using two-bit encoding in a lookup table of k-mer prefixes starting from the PAM site. Since bulges can result in sgRNAs of different lengths and to limit space usage for the index we restrict the prefix length to 16nt (0.5 GB for the human genome). Since 16nt is shorter than a typical sgRNAs, we store all suffixes of each prefix that is present in the genome. This genomic index enables instant lookup of k-mers adjacent to the PAM, followed by an efficient local alignment step to the complete target site (prefix and suffix). If this alignment is within the predefined distance L, a potential OT is identified.

This genomic index allows instant lookup of k-mers, but can not identify OTs at Levenshtein distance greater than 0. To accomplish this, we instead preprocess the sgRNA to generate all possible k-mers within a given Levenshtein distance L. Generating these corresponds to following all paths through an alignment graph (**Figure 2B**, showing L=2) where the number of paths grows exponentially with L (**Supplementary Fig. 4**). Generating these k-mers is therefore a costly operation. However, a key insight is that these paths are identical for all sgRNAs and can therefore be abstracted and reused for all sgRNAs. Exploiting this property, we created a symbolic sgRNA defined not in terms of nucleotides, but by the index of the positions relative to the PAM. We then followed all paths through the alignment graph, performing a k-mer expansion on this symbolic sgRNA to obtain a generalized “symbolic” k-mer outcome termed “symbolic alignments” (**Figure 2C**). Using this precomputed list of symbolic alignments allows us to skip the expansion step for any concrete sgRNA and instead simply substitute the sgRNA nucleotides at their respective positions in the symbolic alignments, thereby obtaining a specific “concrete” k-mer expansion for that sgRNA. Such substitution is efficient and takes a fraction of the time it takes to perform graph traversal for a given sgRNA. When complete information of all OTs is not required, early stopping can be used to further increase speed and abort searches after a maximum number of OTs has been reached.

The combination of these two preprocessing steps (the genomic index and the symbolic alignments) generates a highly efficient sgRNA target search algorithm. When compared to the leading tools SWOffinder, CRISPRitz, and Cas-OFFinder, CHOPOFF is an order of magnitude faster (**Figure 2D**) without losing sensitivity (**Figure 2E**). In fact, most tools do not correctly retrieve the full set of OTs at a given Levenshtein distance, and for our test cases, only CHOPOFF and SWOffinder are capable of correctly retrieving all OTs.

We expanded this algorithm further by allowing ambiguous bases, mismatches and indels to support genomic variation and methods for precision medicine. CHOPOFF-*vcfDB* accomplishes this by extending the genomic index to keep track of all possible ambiguous base combinations, allowing for multiple overlapping variants of any type (insertion/deletion/mismatch). This comes at the cost of a modest decrease in speed, although the process is still significantly faster than the fastest other tool, CRISPRitz (**Figure 2F**).

### Binary Fuse Filters allow for near-instant estimation of off-target events

When a very large number of guides is evaluated and speeds beyond even symbolic alignments are needed, heuristics based on alignment-free pre-filtering of low-quality sgRNAs can produce near-instant results. The recently developed Binary Fuse Filters (BFF) provide fast approximate set membership while using very little memory ^27^. This probabilistic filtering offers a guarantee that it will never miss a sgRNA that is in the set (i.e. no false negatives), but may produce a false positive with low probability. To encode all unique sgRNA and their counts, we created a stack of BFFs where each BFF encodes all sgRNAs with a specific number of OTs (**Figure 3A**). We define sgRNA with 10 or more OTs to be ineligible for editing purposes and store these sgRNAs in one BFF as a final tenth layer. This data structure can be stored using only 503MB of data for the human genome (hg38, UInt16 encoding), as compared to a complete dictionary of all k-mers with 3.1GB. The vast majority of all sgRNAs are unique (95.61%), and their counts follow a long-tailed distribution, enabling a layered database design (**Supplementary Figure 5**).

**Figure 3.**
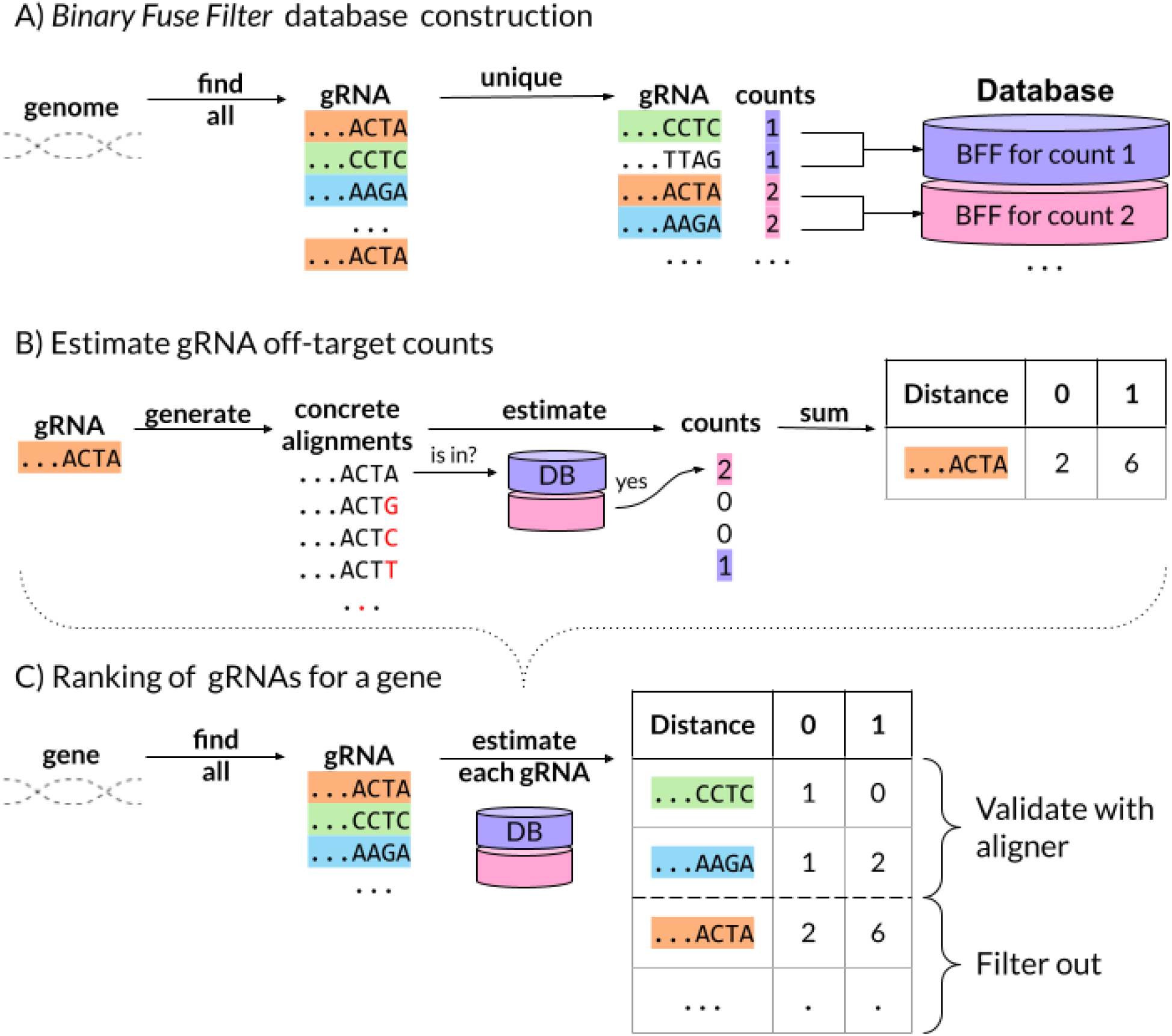
Near-instant alignment-free estimation of off-target counts with CHOPOFF. **A**) Grouping guide sequences by the number of occurrences in the genome allows them to be stored in probabilistic and space-efficient data structures - binary fuse filters (BFF). Each filter stores all guide sequences that appear a specific number of times in the genome (e.g. the first filter contains all guides that appear exactly once). **B)** When a sequence is queried, it is first expanded using symbolic alignments to its concrete alignments. These are sequentially checked for presence in each of the BFFs. The first hit gives an estimate of the number of occurrences for this concrete alignment. The sum of occurrences for all concrete alignments gives an estimate of off-targets. BFFs only make false positive errors at low probability, and there is no risk of false negatives. **C**) For a gene of interest, all sgRNAs can rapidly be filtered for low-quality sgRNAs with high off-target estimates. sgRNAs that pass the filter can be verified with our previously described genomic index approach or with an aligner supporting large distances.

When the OT count of an sgRNA needs to be determined, the guide is first expanded to all possible concrete alignments, and each of these is evaluated in the layered BFFs from low counts to high counts (**Figure 3B**). Due to the probabilistic nature of BFFs, this may result in underestimating the off-target counts for some sgRNAs, but will not reject any sgRNAs that have low off-target counts. Importantly, when sgRNA is reported as off-target free it is guaranteed to be true. Estimating OTs for an sgRNA is achieved through enumerating the occurrence counts for all concrete alignments for a given sgRNA (**Figure 3B,C**). For larger search operations, such as all sgRNA candidates in a gene, this allows for near-instant filtering of all sgRNAs that have a high OT count and restricts exhaustive search to only those sgRNAs that have the potential to be highly specific. For larger servers where space and memory are not an issue, CHOPOFF also supports the use of a traditional dictionary data structure, replacing the BFF, which decreases estimation errors.

## DISCUSSION

We reprocessed several large-scale OT prediction sets and showed that by ignoring insertions and deletions, these methods will miss OT editing. Our results demonstrate that the Levenshtein distance more accurately represents differences between guide RNAs and sites, and that there exist bona fide edited OT sites with a low Levenshtein distance, but high Hamming distance. We also demonstrated that using a Levenshtein distance threshold greater than 3 results in over 99% false positives, and that at this distance sites are being cleaved at negligible rates (**Table 1**). Based on these findings, we suggest that including sites with a distance of 4 or more in OT predictions likely introduces excessive false positives, making them uninformative during guide design. Given the lack of fast and comprehensive tools for OT prediction that account for bulges (**Figure 2D**) and the limitations of current tools in providing complete OT sets (**Figure 2F**), we developed a new method: CHOPOFF.

CHOPOFF introduces methods for OT prediction that are significantly faster and more accurate than the current state-of-the-art methods. Thus, CHOPOFF allows for more sensitive searches and higher quality, particularly relevant for large-scale experiments. It should be noted that CHOPOFF does not provide a general sequence alignment technique; rather, it is a strategy optimized for sgRNAs, as efficiency comes at the expense of requiring separate indices for each PAM. It is also restricted to a limited prefix length that is intractable for mapping longer sequences. However, symbolic alignments introduce a paradigm that can be generalized to speed up other seed-based aligners.

CHOPOFF also has implementations of other proof-of-concept algorithms (see **Supplementary Material** or CHOPOFF documentation), such as Vantage-point tree ^28^, pigeonhole principle ^29^, 01*0 seed ^30^, which were implemented as part of our research into the most performant solution. The methods are open source (**Table 2**), implemented in Julia and compatible with crisprVerse ^31^ through the crisprCHOPOFF R package. All CHOPOFF methods have above 95% test coverage, ensuring robustness and high quality of the software.

**Table 2.** CHOPOFF links presented in the paper.

| Repository | Description |
| --- | --- |
| https://crisprtools.org/chopoff | CHOPOFF with web interface |
| https://github.com/JokingHero/CHOPOFF.jl<br>DOI: 10.5281/zenodo.14039564 | CHOPOFF software, Julia package |
| https://jokinghero.github.io/CHOPOFF.jl/ | CHOPOFF documentation |
| https://github.com/JokingHero/crisprCHOPOFF<br>DOI: 10.5281/zenodo.14039583 | R package with CHOPOFF interface |
| https://git.app.uib.no/valenlab/chopoff-benchmark | Benchmark code and results |
| https://git.app.uib.no/valenlab/offtarget_reanalysis | Re-analysis of available datasets |

Taken together, CHOPOFF is a fast and sensitive method that addresses the main challenges associated with predicting potential off-target sites for CRISPR/Cas assays. Importantly, CHOPOFF supports the prediction of OT in the context of genomic variation, a crucial requirement for the use of CRISPR/Cas in personalized medicine.

## MATERIALS AND METHODS

### Benchmarks

For the benchmark, all tools used the same reference for OT search, the human genome (GENCODE hg38 v34, Dec 2013), which each tool preprocessed to its own format. Database construction is not part of the benchmark. The CHOPOFF prefixHashDB method was run with early stopping conditions set to a maximum of 10^6 OTs for each sgRNA at each distance, ensuring it found all OTs and did not utilize its early stopping condition. We searched with the same 61 sgRNAs for each tool and reported the mean running time. For the purpose of the analysis, we allowed a Levenshtein distance up to 4, which ensured inclusion of bulges and mismatches. Cas-Offinder was installed from GitHub (v 3.0.0, supporting bulges) and run on a GPU (TigerLake-H GT1, faster than 15 threads on CPUs), while CRISPRitz (v 2.6.6) was installed through conda and run with 15 threads, and CHOPOFF (v1.0.0) was also run with 15 threads. Benchmarks were run on a regular laptop machine with 32GB RAM and 16 processors, 11th Gen Intel(R) Core(TM) i7-11850H @ 2.50GHz. The code link is available at **Table 2** below.

### Output comparison of off-target searches

We restricted our comparison of the different tools to results at Levenshtein distance L<=3 because above this, any given predicted site has close to zero percent probability of being cut, and if it is indeed cut, the efficiency is typically extremely low (**Figure 2A**). In addition, CRISPRitz produces highly redundant output showing many alignments for the same OT sites, and for distance 4, its output of 160GB is too large to parse. Each program outputs OTs’ positions differently, and we therefore gave them the benefit of the doubt, allowing for 5bp imprecision in reporting the OT position. All of CHOPOFF’s algorithms return exactly the same result, in the same format, and SWOffinder reports identical OT sites to CHOPOFF. Both CRISPRitz and Cas-OFFinder do not find any OT sites that are not also found by CHOPOFF and SWOffinder. However, CRISPRitz does not find 3058 sites, and Cas-OFFinder does not find an additional 7860 OTs. We verified these OTs through independent string searching, showing that the sites found exclusively by CHOPOFF and SWOffinder are indeed correct, and other tools missed them. In the repository (chopoff-benchmark), we provide the file ‘compare_software_results.html’, which contains examples of these OT sites that are not detected. We notice that all OTs not reported by CRISPRitz and Cas-OFFinder are for distances 2 and 3, they come from a variety of sgRNAs and from different sites. While we have not further explored the details of their omission, we note that CRISPRitz missed sites are always in the form of a combination of bulges between the reference and sgRNA.

### Cell-free screening

To simulate potential off-target editing, 27 different guide RNAs (gRNAs) were designed and ordered through Integrated DNA Technologies (IDT). These gRNAs were designed to contain different mismatches, insertions and deletions compared to the wild-type (WT) sequence. crRNAs were annealed to tracrRNA to create gRNAs following the protocol recommended by IDT. gRNA sequences are listed in Supplementary Methods. A segment of the ADA2 gene was used as the WT template and was produced through nested PCRs. The resulting fragment was 946 bp long, where the Cas9 would cut asymmetrically to create differently sized fragments for ease of visualization in a gel. Primer sequences are listed in Supplementary Methods. Guide-it™ sgRNA Screening Kit (Takara Bio, Cat#632639) was used according to the manufacturer’s protocol to test the 27 guides and their cutting potential when digested with the template DNA. The resulting DNA fragments were visualised in a 2% agarose gel.

## DATA AVAILABILITY

## SUPPLEMENTARY DATA

**Supplementary Data are available online**.

## AUTHOR CONTRIBUTIONS

Kornel Labun: Conceptualization, Implementation, Validation, Writing. Oline Rio and Anna Zofia Komisarczuk: Validation in the lab, Writing. Håkon Tjeldnes and Michał Swirski: R package and crisprVerse integration, Writing. Emma Haapaniemi: Writing, and Eivind Valen: Conceptualization and Writing.

