## Supplementary Material for "CHOPOFF: symbolic alignments enable fast and sensitive CRISPR off-target detection"

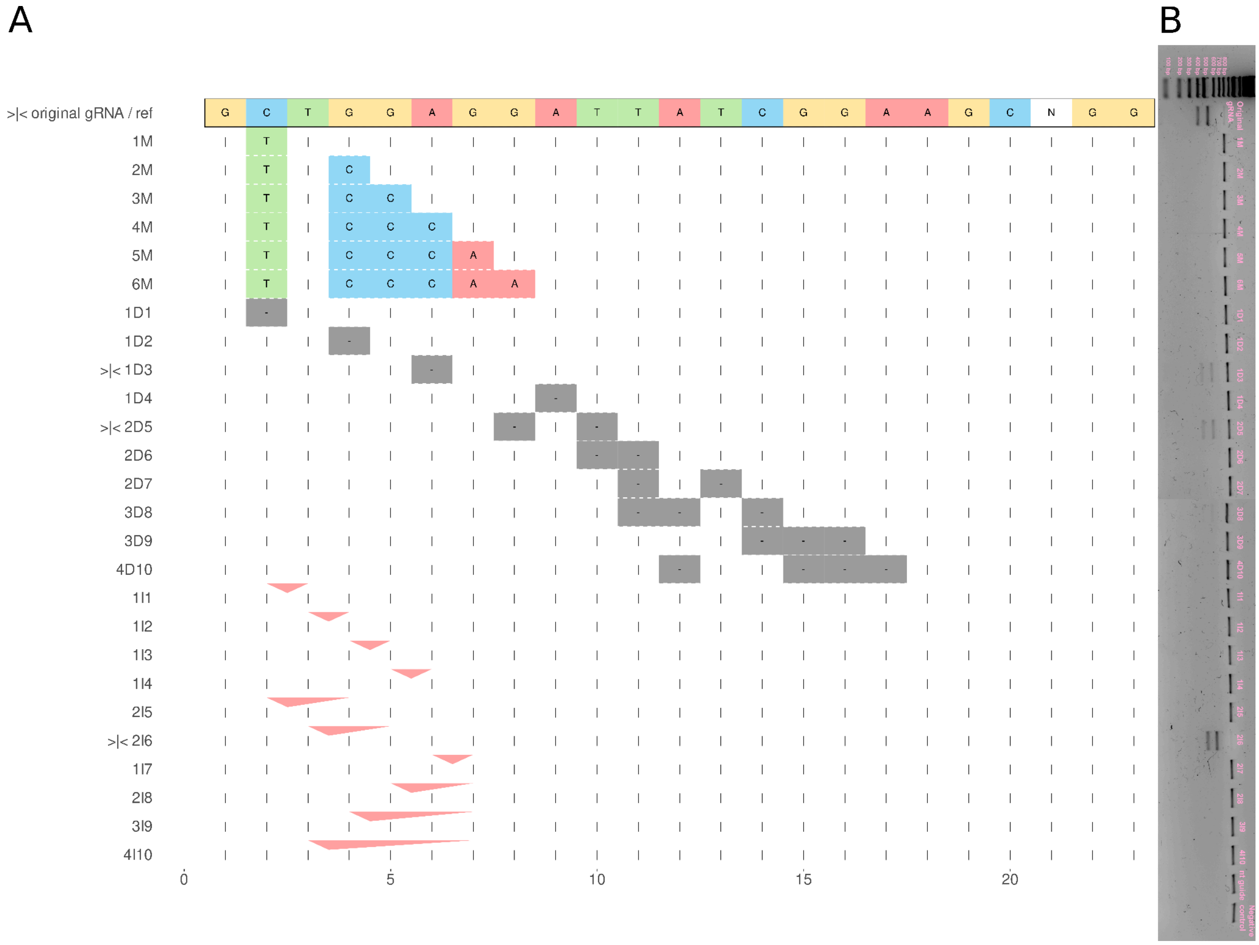


**Supplementary Figure 1**. **A**) Design of sgRNA-OT pairs with variable distances and mismatches, deletions (grey dash) and insertions (red triangle, bottom points to the position of the gap) that relies on the same reference and different sgRNAs to simulate multiple OTs. OTs which show cutting are marked with “>|<”. **B**) Cell-free screening with gel electrophoresis. Original sgRNA, 1D3, 2D5 and 2I6 exhibit cutting of the DNA.

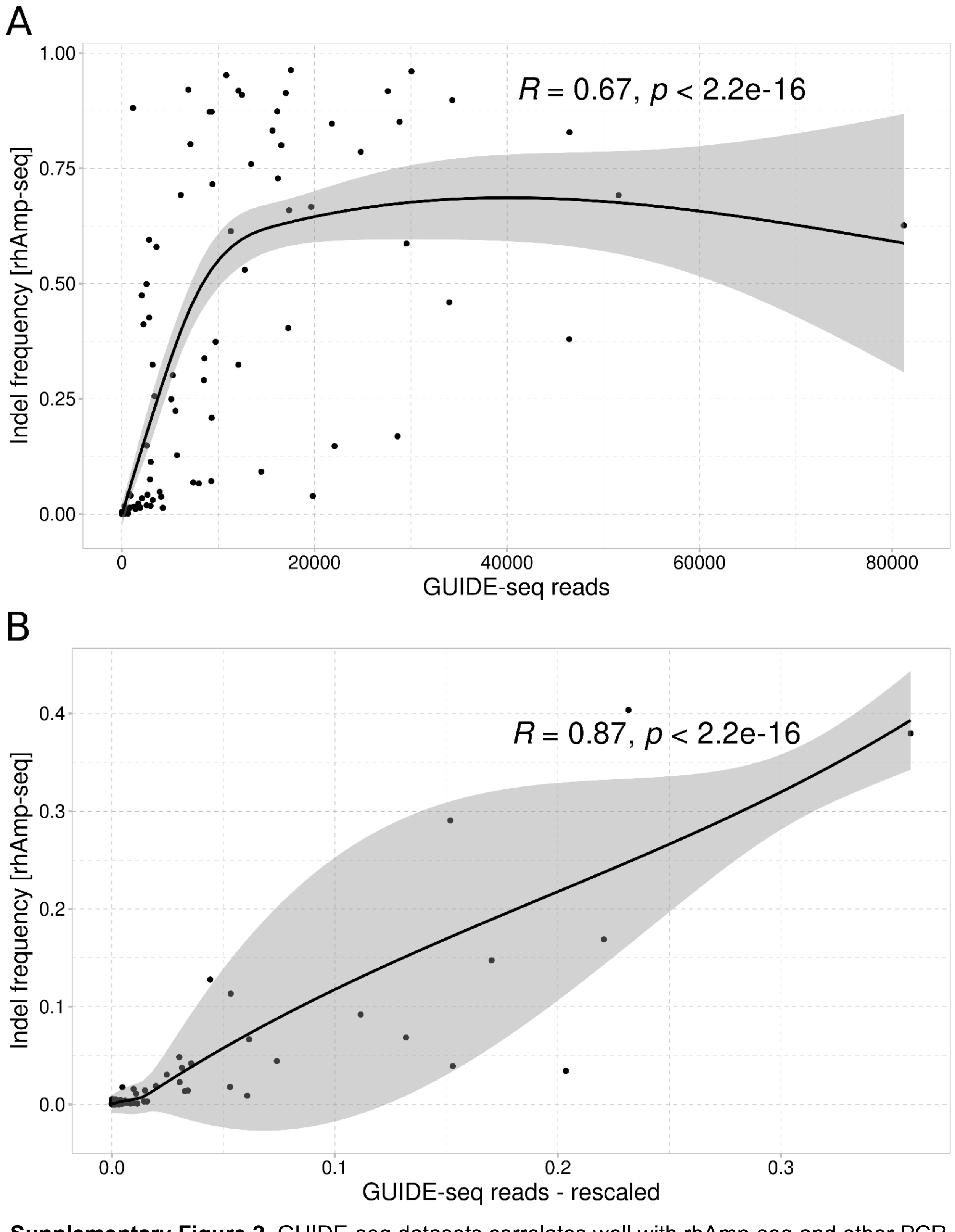


**Supplementary Figure 2**. GUIDE-seq datasets correlates well with rhAmp-seq and other PCR sequencing validation (**A**), they can be also rescaled using on-target editing efficiency (**B**) to approximate off-target editing efficiencies.

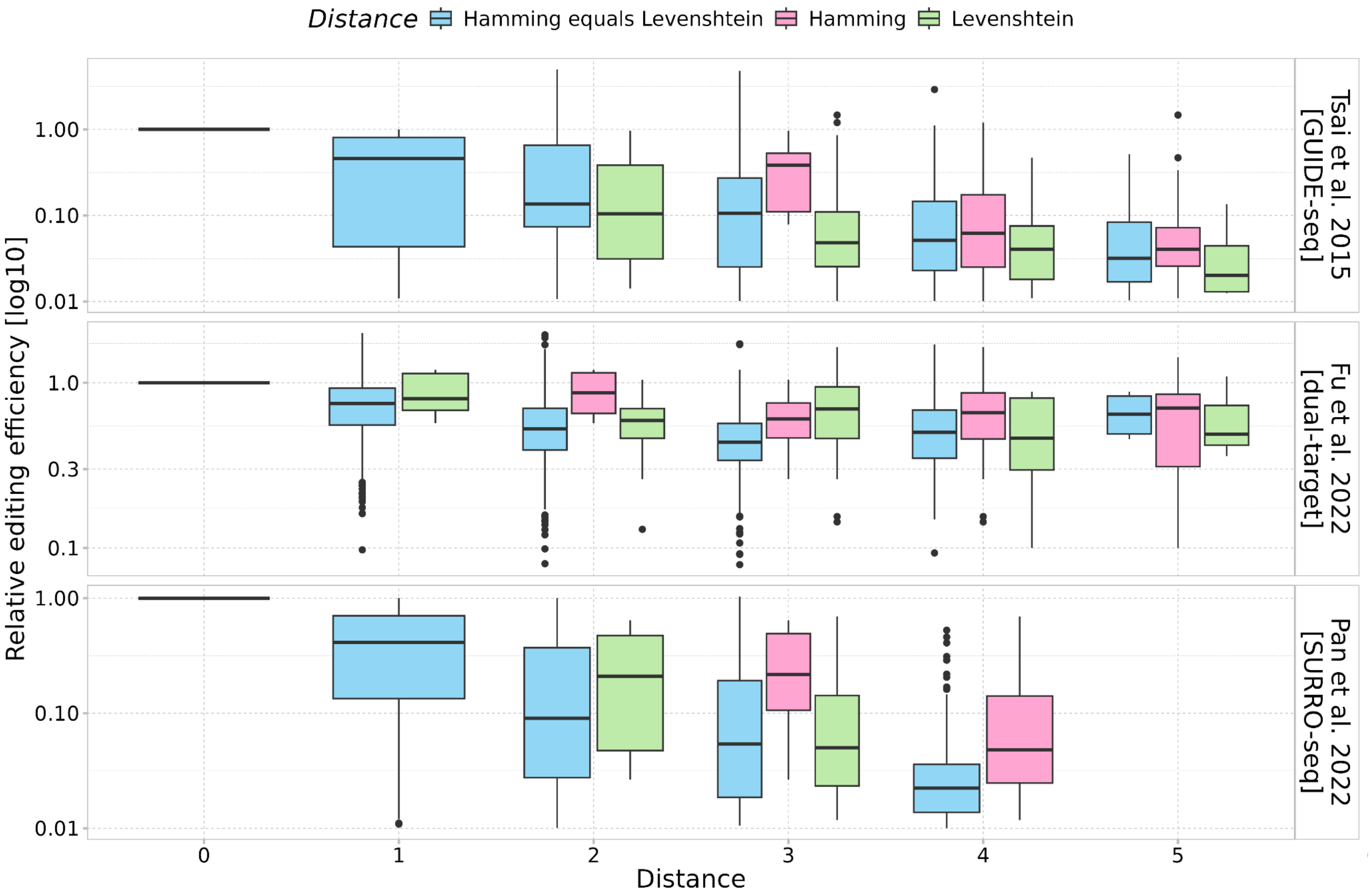


**Supplementary Figure 3**. Comparison of Hamming (red) and Levenshtein (green) metrics for off-target alignment when the metrics disagree versus when they agree (blue). Three different biological datasets are shown (GUIDE-seq [1,2], dual-target [3], SURRO-seq [4]). Off-target activity is normalized to the editing efficiency of the on-target site. The Levenshtein metric (green) more closely resembles the ground-truth relationship (blue boxes) than the Hamming metric (red), indicating it is a better predictor of editing efficiency for sites where the metrics disagree.

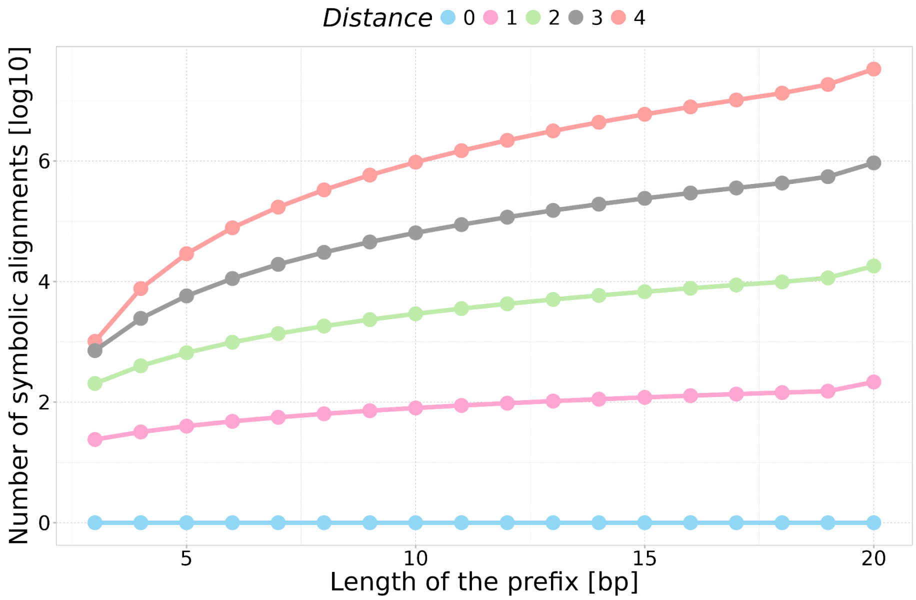


**Supplementary Figure 4**. Number of symbolic alignment paths in relation to length of the prefix.

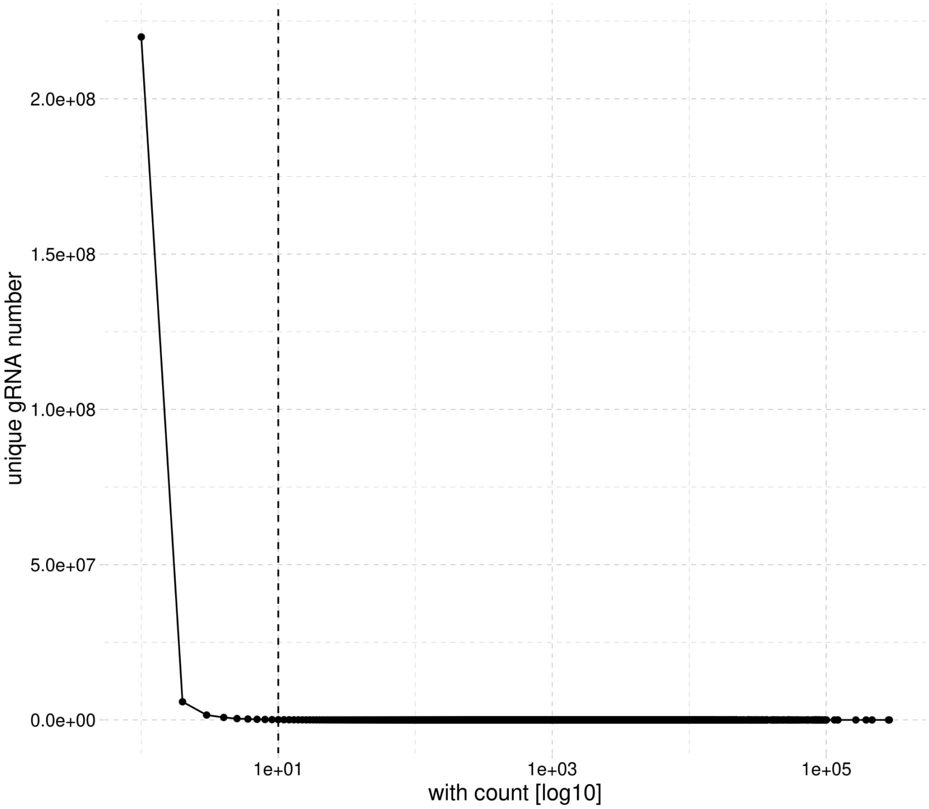


**Supplementary Figure 5**. Distribution of 230M unique sgRNA for hg34 with their respective count. The cutoff is a value count of 10 which captures 95.61% of all gRNAs. The vast majority of the gRNAs are unique, but there exist outliers with thousands of replications across the genome.

### Detailed description of the CHOPOFF algorithms

CHOPOFF contains many algorithms and methods for OT searching that can be used interchangeably. They were implemented during our search for a more performant solution, each of them has its own strengths and weaknesses. For current and up to date documentation of all CHOPOFF methods, functions and classes please visit the official documentation page (https://jokinghero.github.io/CHOPOFF.jl/). Comparison of speed for all of CHOPOFF algorithms is shown in **Supplementary Figure 6**.

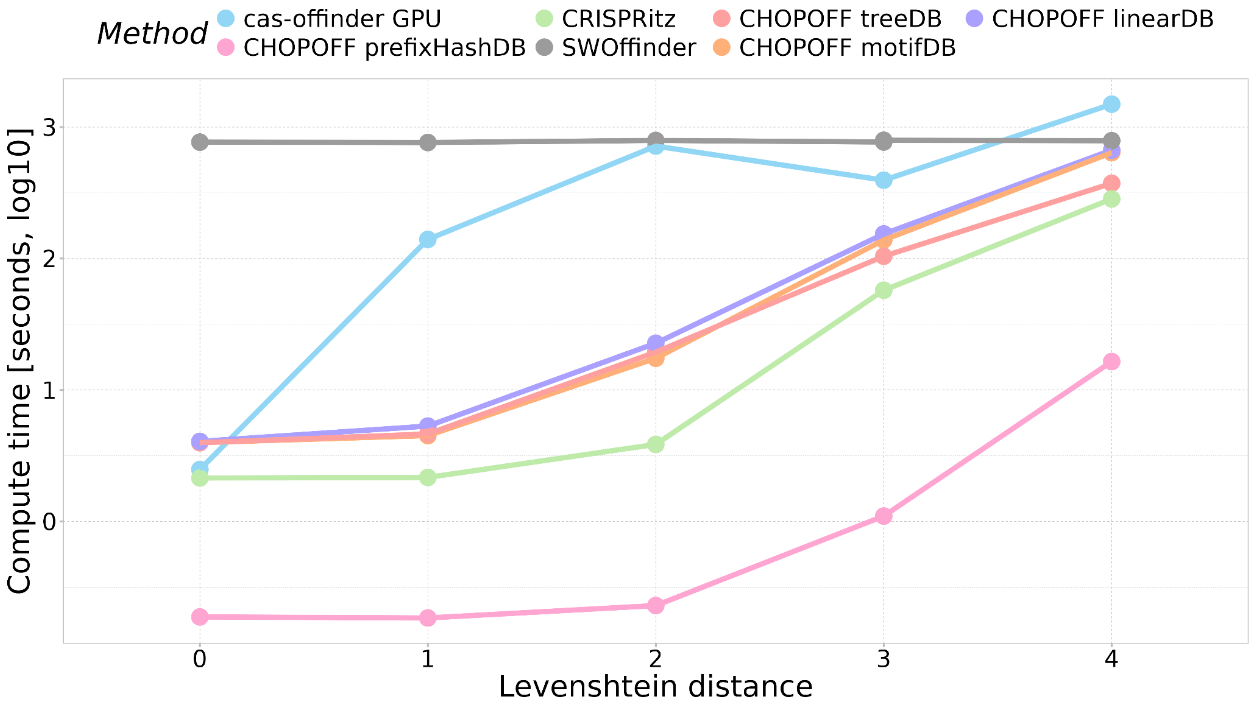


**Supplementary Figure 6**. Benchmark of main CHOPOFF methods. All CHOPOFF methods shown here find all off-targets, each offers a unique approach described below.

##### Prefix-Suffix Methods for exact off-target search with arbitrary Levenshtein distance

Prefix-Suffix methods are based on the idea of splitting the large database of guides into smaller chunks that can be processed independently and therefore enable optimizations such as filtering, partial-alignment, compression and parallelization. Prefix-suffix methods therefore build a database of all potential OT sites which are grouped by their prefix; prefix length is defined by the user during the build step. For example, searching up to a distance of four, our recommended prefix length is eight, but for computers with less RAM or smaller disk space use a smaller prefix.

All methods that use databases for finding OTs suffer from potential overlap conflict. Databases contain all possible OTs for a given PAM sequence, for example for Cas9 NGG and genomic stretch of guanines results in many OTs that each might have valid alignment towards sgRNA. However, naturally, the loci with smallest distance will be preferentially recognized by Cas9 and cut with higher editing efficiency. Looking at all possible OTs hinders the comparison of how many loci on the genome is actually in danger of being cut, therefore CHOPOFF includes a method (see *filter_overlapping* documentation) for filtering out these overlap conflicted OTs. Furthermore, CHOPOFF contains a summarization (see *summarize_offtargets* documentation) method which can be used to rank sgRNAs by their number of OTs.

**linearDB is a method for exact off-target search with the support for early stopping**

linearDB is the simplest implementation out of all CHOPOFF algorithms, it partitions all OT sites by their prefix and stores them in the database. It follows a similar design as CRISPR-SE software, but is expanded to allow bulges instead of mismatches only[5]. During the search step it initially aligns to the prefixes for each of the sgRNAs in parallelized fashion. It is utilizing an extensively optimized needleman-wunsch algorithm with bounded distance for alignment between sgRNA and OT. This alignment is optimized for CRISPR, meaning it is anchored at the Protospacer Adjacent Motif (PAM) and extended towards the 3' end. When all bases of sgRNA are aligned and not full OT sequence is used (e.g. when sgRNA has bulges) the leftovers of OT are not being counted towards the alignment, which is different to the standard edit/levenshtein distance alignment algorithm between two sequences. During the initial search step, if partial alignment between sgRNA and prefix is possible (within specified distance) alignment will be extended for each of the OT suffixes contained in that prefix. All OTs are then reported in the output file, together with alignment, and location of each OT. This method incorporates bulges, ambiguous bases and early stopping,  due to lack of additional filtering it sacrifices some speed. Thanks to extensive testing and simplicity of this method it became our gold standard.

**motifDB is faster than linearDB due to filtering with pigeon-hole principle or with lossless seed**

motifDB follows the same principles as linearDB algorithm, but it adds a filtering layer on top of it. The purpose of this complication is to decrease the time of OTs search further and allow us to experiment with different filtering rules. Currently, no early stopping is implemented, as opposed to linearDB, but it could be added in the future. motifDB keeps track of which spaced k-mers of suffixes belong to which OT (**Supplementary Figure 7**). Although the spaced k-mer order is lost, it is still possible to manipulate the requirements of the number of k-mers that must match between OT and sgRNA given the number of errors already spent on the alignment between prefix and sgRNA. With the current implementation it is possible to adjust all parameters in this filtering scheme and create filters similar in principle to pigeon hole[6] or 01*0 lossless seed[7].

For example, we are interested in finding OTs within levenshtein distance 4 (d) for Cas9 sgRNAs which are 20bp long. We skip prefix length of 7bp, and are left with 13bp of suffixes. Suffixes can be split into skipping 3-mer (r) of size 4, 1bp will be left unused. Now imagine that during searching we encounter a prefix where initial alignment was of distance 3 (m) and adjustment parameter is 0 (a). We are obliged to find at least r - (d - m + a) skip-mers inside the OTs, which in this concrete example evaluates to 3 - (4 - 3 + 0) = 2. Therefore we can filter out all potential suffixes that do not have at least 2 of our 3-mers matching with the sgRNA.

There exist also another approach which builds on the idea that it might be more efficient to find at least two kmers of smaller size (named 01*0 lossless seed) rather than one larger k-mer (pigeon hole principle). You can use the adjustment parameter for that during the search step. Be sure to understand implications of using motifDB filtering schemes as using wrong parameters might result in leaky filtering in relation to the assumed distance during the search step.

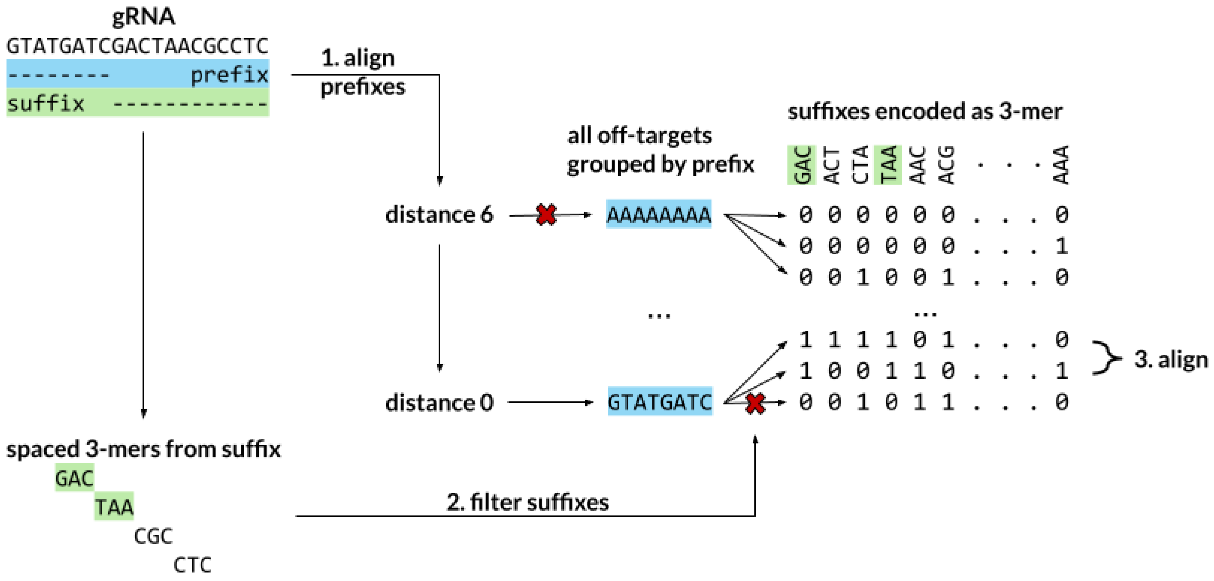


**Supplementary Fig 7.** motifDB filtering scheme operates based on spaced k-mers of the suffix part of the potential off-targets. Here, green 3-mers represent an off-target which enables further filtering (red x) for the prefixes that pass standard linearDB filtering in the first step.

**treeDB uses vantage-point tree for alternative searching scheme**

treeDB still partitions the data into smaller bins based on the prefixes, but it uses the vantage-point tree[8] for its queries against suffixes. Vantage-point tree is using the fact that levenshtein distance belongs to the metric space and therefore we can use the triangle inequality principle between: current vantage-point tree node, sgRNA, and the OT. This allows us to prune the nodes of the tree that are in the inside or the outside (see **Supplementary Figure 8**). Same CRISPR specialized alignment algorithm is used for treeDB as for linearDB and motifDB, however the treeDB inequality principle introduces an inconvenience. Due to the gaps in sgRNA we have to include extensions for vantage-point (VP) and Node alignment (see **Supplementary Figure 9**). treeDB contains a unique filtering scheme that is particularly useful for longer sequences and will prove faster than motifDB for alternative Cas effectors.

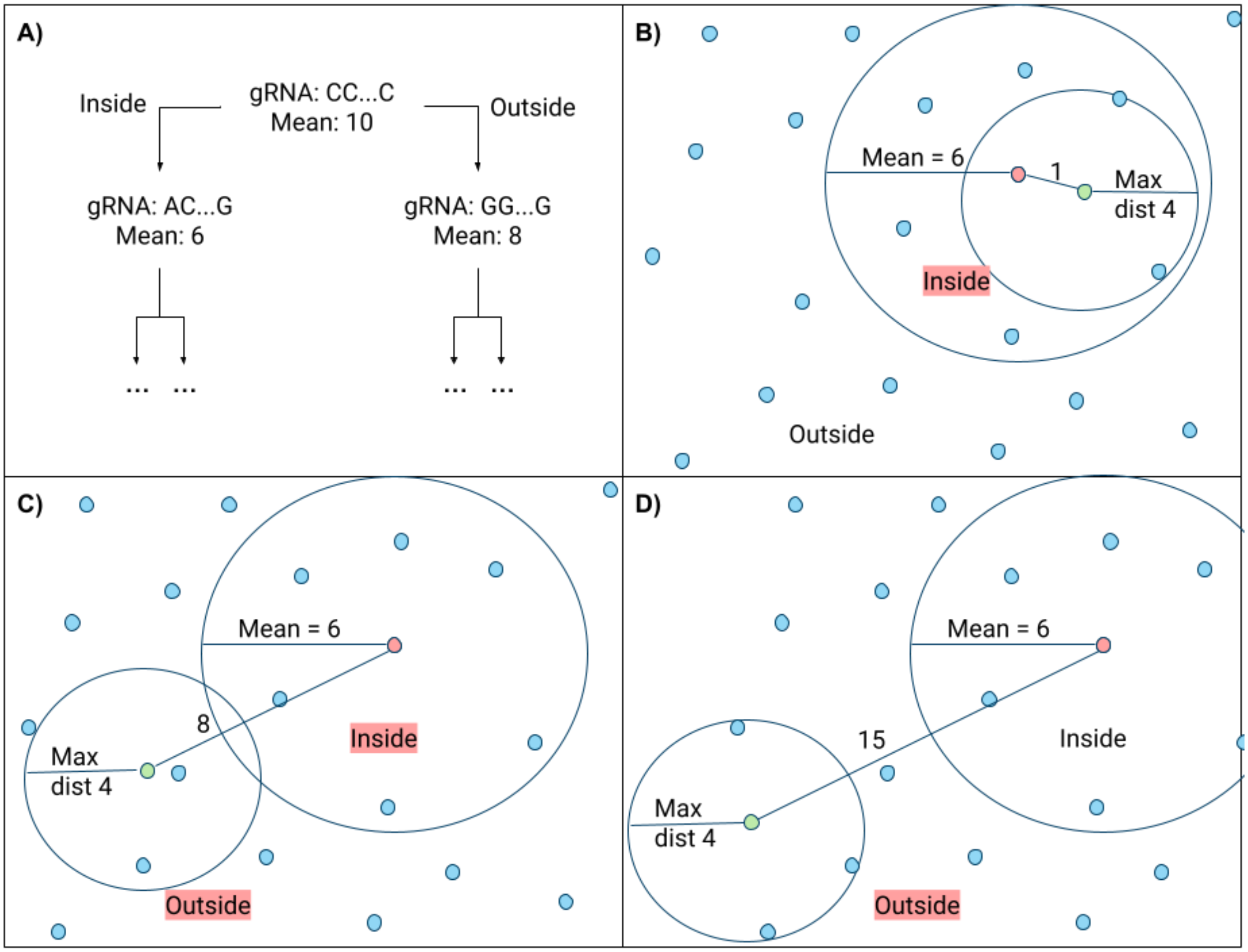


**Supplementary Figure 8**. treeDB is using vantage-point (VP) tree for off-target search. **a)** Each VP contains its sgRNA sequence and mean of distances to all of its nodes. *Inside* subtree contains all sgRNA of distance less or equal to the mean and the *Outside* subtree contains larger distances than the mean. **b)** During off-target search if sgRNA (green dot) has small distance towards the VP (red dot) it is possible that only the *Inside* subtree has to be searched for off-targets. **c)** Worse case scenario, where both *Inside* and *Outside* have to be searched due to maximal distance being too large in relation to the mean of that VP. **d)** Distance between sgRNA and VP is large enough that only the *Outside* subtree has to be searched.

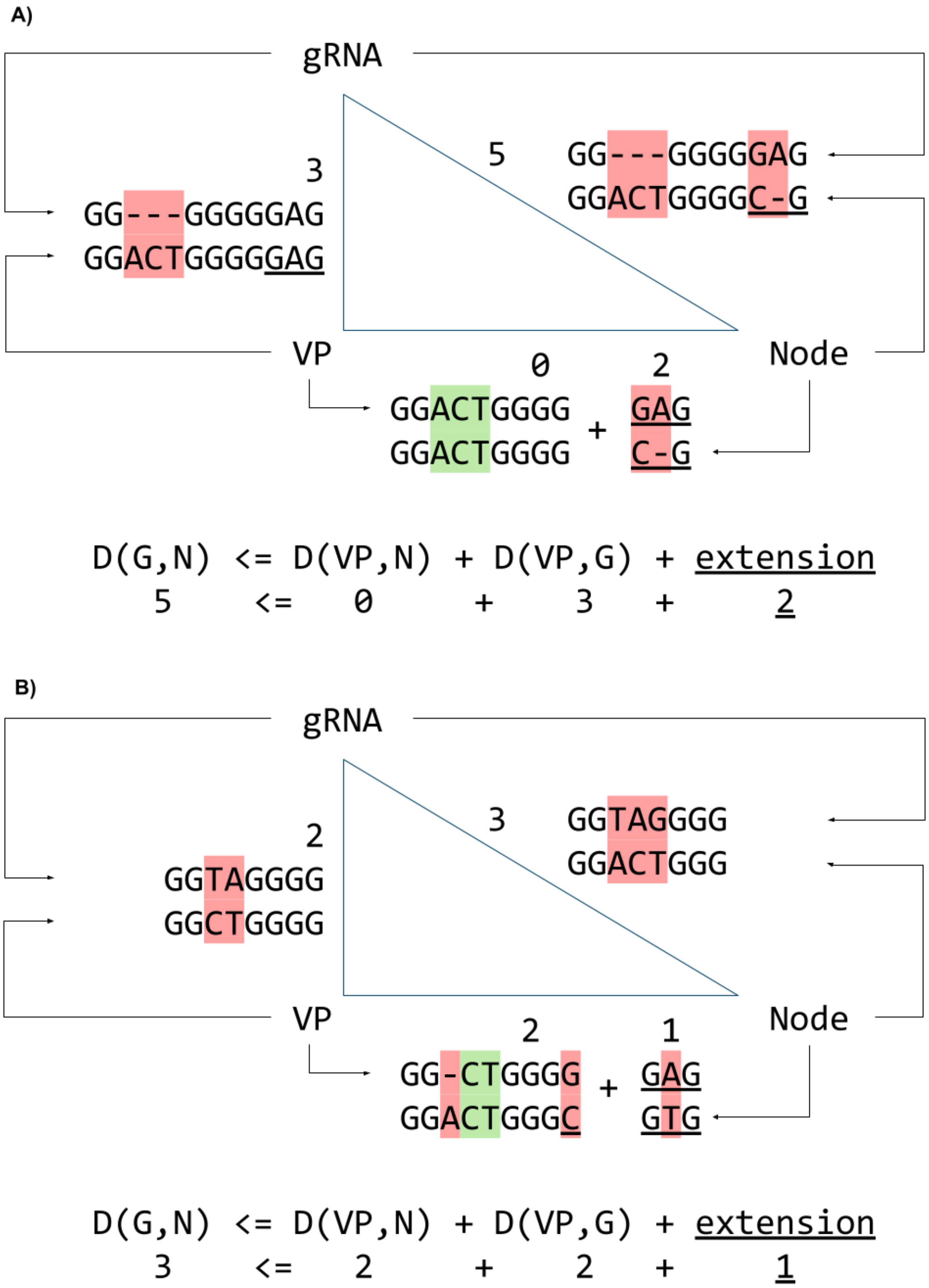


**Supplementary Figure 9**. Extension (underscored) calculations in treeDB for CRISPR specific alignment between sgRNA, vantage-point (VP) and Node when using triangle inequality rule. Example **a)** shows the need to include extensions for vantage-point and Node alignment (due to gaps in sgRNA), while example **b)** shows the case where no gaps are in the sgRNA alignment

**fmiDB is using FM-index for traditional search using enumeration**

Traditionally, FM-index and its derivatives are used for genomic sequence alignment and we have also explored this avenue. We present a method that is relatively fast, but only for smaller distances (up to distance of 2) and becomes impractically slower for larger distances. fmiDB does not suffer from overlapping OTs problems to the same extent as other methods. fmiDB search is optimized to firstly enumerate all possible OTs for each sgRNA sequence in an efficient manner by precomputing all the graph paths (**Figure 1B**). For each possible OT subsequence of sgRNA we then use FM-index for the search of the genomic locations.

**vcfDB allows to search off-targets overlapping SNPs, indels and other variants**

*vcfDB* algorithm is based on a similar principle as *prefixHashDB*. *vcfDB* keeps track of the hashes for each possible ambiguous bases combination. CRISPRitz mode of operation where SNPs are first inserted into the genome in their respective loci, are not able to accommodate insertions and deletion variants, because these would shift the annotation of all upstream loci on that chromosome. *vcfDB* builds a database of all possible OTs with variants (including mismatches/deletions/insertions) and even allows for overlapping variants. *vcfDB* keeps track of the original OTs location and additional annotations for each variant. *vcfDB* method is also used when the definition of CRISPR effector in *prefixHashDB* allows ambiguous bases, because standard pipeline does not keep track of the ambiguous hashes. *linearDB*, *motifDB*, *treeDB* are compatible with ambiguity in the CRISPR effector on their own, because they don't rely on hashing.
